# A recombinant *dilp2GS-rpr* donor line for adult-inducible IPC ablation across *Drosophila* genetic backgrounds

**DOI:** 10.64898/2026.06.17.733056

**Authors:** Yuyan Chen, Yulin Bai, Xuan Zhuang

**Author notes:** **Corresponding author:** Correspondence to Xuan Zhuang.

## Abstract

Genetic-background studies require defined perturbations that can be crossed reproducibly into many recipient backgrounds. We generated a *Drosophila dilp2*GS*-rpr* donor line for adult-inducible ablation of insulin-producing cells (IPCs), which secrete insulin-like peptides and provide a tractable model of insulin-deficient metabolic physiology. This line carries *dilp2*-GeneSwitch-GAL4 and UAS-*reaper* in cis on the same second chromosome homolog over a balancer. PCR genotyping and sequencing confirmed both transgenic elements in the candidate recombinant line. RU486 induction reduced *dilp2* mRNA expression, supporting partial IPC ablation. Treatment-duration testing identified 8 days of RU486 as sufficient to increase whole-body glucose in the *dilp2*GS*-rpr* line but not in the background-matched control; food intake did not differ between RU486- and vehicle-treated flies. Across metabolic assays, whole-body glucose showed the clearest RU486- and line-dependent phenotype. This validated *dilp2*GS*-rpr* line enables testing how recipient genetic backgrounds modify inducible IPC/DILP metabolic phenotypes and provides a framework for similar linked donor-line resources.

## Introduction

Genetic-background studies are most scalable when a defined perturbation can be packaged into a standardized, traceable donor genotype and crossed reproducibly into many recipient backgrounds. *Drosophila melanogaster* is a powerful genetic model for studying how defined perturbations interact with genetic background to shape complex traits including metabolism, reproduction, and lifespan. The *Drosophila* Genetic Reference Panel (DGRP) provides a well-characterized resource for mapping the effects of natural genetic variation on complex traits (Mackay et al., 2012). This donor-line strategy can be applied broadly to GAL4/UAS-based perturbations; here, we apply it to the insulin-producing cell (IPC)–DILP system as a model for adult metabolic modifier studies. Although flies lack a pancreas, brain insulin-producing cells (IPCs) secrete DILP2, DILP3, and DILP5 and function as endocrine regulators analogous to mammalian pancreatic β-cells. Altered IPC function or reduced insulin-like signaling affects carbohydrate and lipid homeostasis, growth, reproduction, stress responses, and lifespan (Broughton et al., 2005; Rulifson et al., 2002), making the IPC–DILP axis useful for modeling insulin-deficient states physiology and mapping genetic modifiers of IPC/DILP-dependent metabolic traits.

Existing IPC and DILP perturbation models have provided important insights, but systematic analysis of adult phenotypes and their modifiers across many genetic backgrounds requires additional features. Temporal control is needed for adult endpoints because IPC/DILP signaling acts through the insulin/insulin-like signaling pathway during development; early IPC or DILP perturbation can alter growth, developmental timing, viability, and adult physiology(Rulifson et al., 2002; Shingleton et al., 2005; Zhang et al., 2009). Adult induction therefore reduces developmental confounding. Modifier mapping across diverse genetic backgrounds is facilitated when perturbing elements can be inherited together as a single genetic unit. The GeneSwitch-GAL4 system enables RU486-dependent activation of UAS-linked transgenes, and *dilp2*-GeneSwitch/UAS-*reaper* has been used previously to induce adult IPC ablation or functional knockdown and to study glucose homeostasis, energy storage, fecundity, and lifespan(Haselton et al., 2010). Inducible systems also require matched controls to distinguish target-specific induction from effects caused by the inducer, basal transgene activity, or the genetic background itself, as illustrated by prior *dilp2*-GS/UAS-*rpr* studies that included uninduced or vehicle-treated controls(Serbus et al., 2015; Tanabe et al., 2017). Thus, the gap addressed here is not the absence of IPC-ablation models, but the need for an optimized donor-line resource that makes this perturbation portable across genetic backgrounds.

In this study, we generated the *dilp2*GS*-rpr* donor line, a genetically portable, temporally inducible IPC-ablation resource in which *dilp2*-GS and UAS-*rpr* are linked in cis on a single recombinant chromosome maintained over a balancer. This strategy builds on previous DGRP modifier studies in which standardized donor or tester stocks carrying linked Gal4/UAS perturbation elements were crossed to DGRP lines to map natural genetic modifiers of defined phenotypes (He et al., 2014; Palu et al., 2022; Park et al., 2014). After establishing the recombinant chromosome, we backcrossed the donor line to a background-matched control line and validated the model using molecular and metabolic readouts. We used matched *dilp2*GS*-rpr* donor-line and control groups with and without RU486 treatment to separate IPC-ablation-specific effects from RU486-dependent effects and baseline differences associated with the recombinant chromosome. This resource provides a linked, balanced recombinant donor chromosome for mapping modifiers of adult IPC/DILP phenotypes, with whole-body glucose serving as the primary validated metabolic readout. More broadly, the construction and validation workflow provides a template for generating similar linked donor-line resources from other compatible transgenic perturbation systems.

## Materials and Methods

### Fly stocks and husbandry

The *dilp2*-GeneSwitch-GAL4/*CyO* driver line was obtained from Dr. Heinrich Jasper’s lab at the Buck Institute. The UAS-*reaper* line (Bloomington *Drosophila* Stock Center, stock #5824) and the *sna[Sco]*/*CyO* line (Bloomington *Drosophila* Stock Center, stock #8578), used as a matched background control, were obtained from the Bloomington *Drosophila* Stock Center (**Supplementary Table S1)**. All fly stocks were maintained at 25□ °C in a temperature-controlled incubator with 60–65% humidity under a 12 h:12 h light/dark cycle. Flies were reared on standard cornmeal–yeast–molasses medium during stock maintenance and development (**Supplementary Table S2 Standard Maintenance Diet**).

### Construction of the *dilp2*GS-*rpr* recombinant donor line

To generate an inducible IPC-ablation line, we recombined the *dilp2*GS-GAL4 driver and UAS-*rpr* responder onto the same second chromosome. The UAS-*rpr* line (carrying two copies of *mini-white*) was first crossed to the *dilp2*GS-GAL4/*CyO* driver line (carrying one copy of *mini-white*). F1 females harboring the two transgenes on opposite second-chromosome homologs were then crossed to *sna[Sco]/CyO* males to recover recombinant chromosomes generated during female meiosis **(Figure 1A)**. Progeny were screened based on eye color and wing morphology. Mini-white-associated eye pigmentation and the *CyO*-associated Curly wing phenotype were used to enrich for candidate recombinant carriers, but eye color alone was not treated as definitive genotype confirmation (**Figure 1B**). Among approximately 200 screened flies, a candidate recombinant male with orange eyes was identified. This male also exhibited the *CyO*-associated Curly wing phenotype, consistent with recovery of a candidate recombinant chromosome balanced over *CyO*. The recombinant chromosome was subsequently crossed into a *sna[Sco]/CyO* background, and the resulting line was backcrossed for seven generations to generate a background-matched donor stock (**Figure 1A**). We refer to the established balanced stock as the *dilp2*GS*-rpr* donor line.

**Figure 1.**
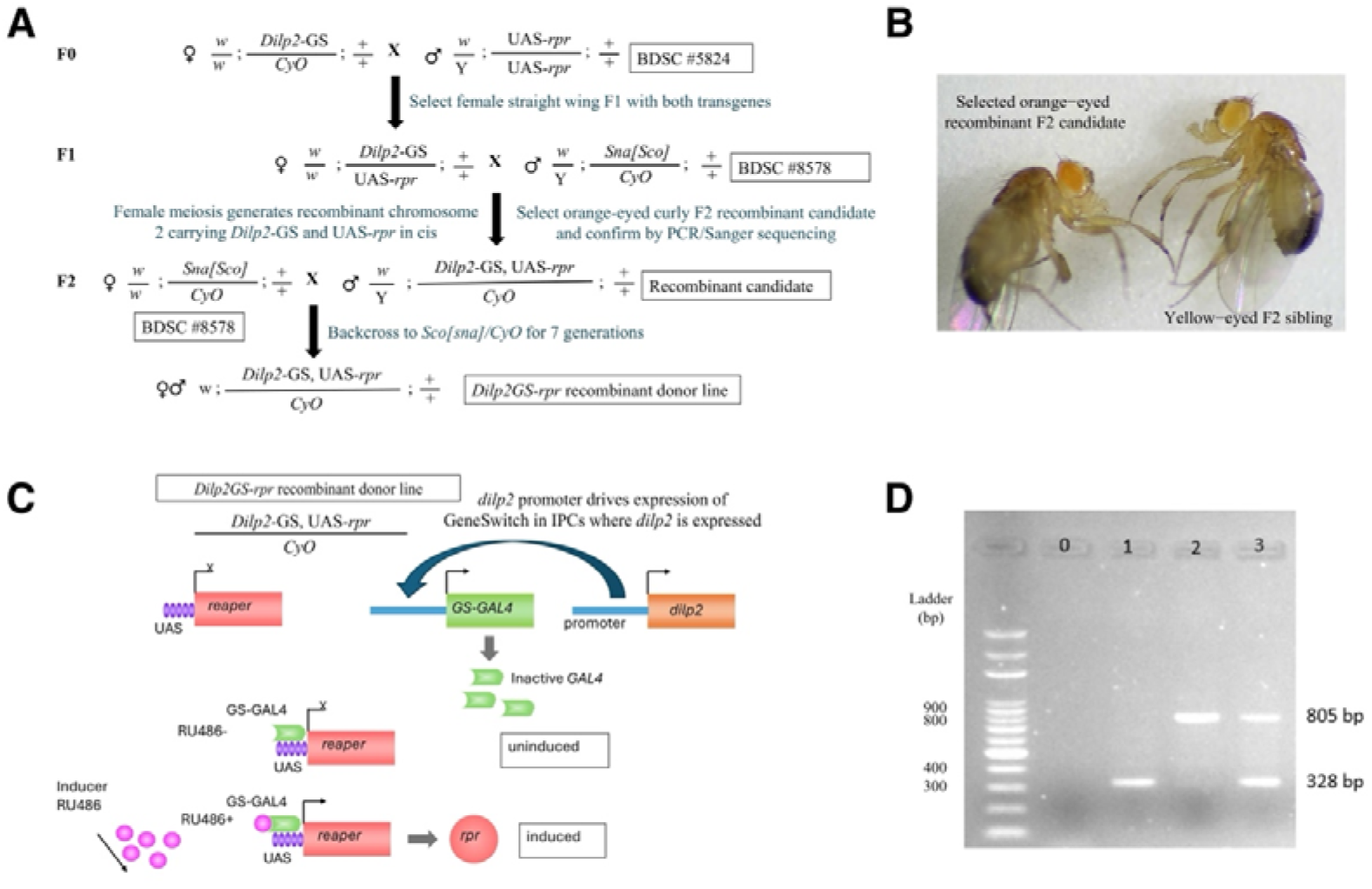
Construction and genetic validation of the inducible *dilp2*GS*-rpr* recombinant line. (A) Crossing scheme used to generate the recombinant donor chromosome. The UAS*-rpr* responder line was crossed to the *dilp2*GS*-GAL4/CyO* driver line. F1 females carrying *dilp2*GS*-GAL4* and UAS*-rpr* on homologous second chromosomes were crossed to *sna[Sco]*/*CyO* males to recover recombinant chromosomes generated during female meiosis. Candidate F2 flies carrying a recombinant second chromosome over *CyO* were selected and the resulting donor chromosome was backcrossed to the matched-control background for seven generations. (B) Representative F2 flies from the recombinant-screening cross. The orange-eyed fly was selected as a candidate *dilp2*GS-rpr recombinant carrier, whereas a lighter/yellow-eyed F2 sibling is shown for comparison. Eye color was used as a screening marker for candidate recombinant flies. (C) Schematic of RU486-inducible IPC ablation in the *dilp2*GS*-rpr* donor line. The *dilp2* promoter drives GeneSwitch-GAL4 expression in adult insulin-producing cells. In the absence of RU486, GeneSwitch-GAL4 is expected to remain inactive and *UAS-rpr* is not induced. After RU486 treatment, activated GeneSwitch-GAL4 drives *rpr* expression from the UAS responder, leading to Reaper-mediated IPC ablation. (D) PCR genotyping of parental and candidate donor lines. Lane 0, no-template genomic DNA control; lane 1, *dilp2*GS*-*GAL4 parental line; lane 2, UAS*-rpr* parental line; lane 3, candidate *dilp2*GS*-rpr* recombinant donor line. Expected PCR products are 328 bp for *dilp2*GS*-*GAL4 and 805 bp for UAS*-rpr*. Detection of both products in the candidate donor line supports the presence of both transgenic elements; amplified products were further verified by Sanger sequencing.

Adult-specific IPC ablation is expected to be induced using the inducible GeneSwitch system (Osterwalder et al., 2001). In this system, the *dilp2* promoter drives GeneSwitch-GAL4 expressions specifically in IPCs. Upon administration of RU486, GeneSwitch-GAL4 is activated and induces expression of the pro-apoptotic gene *reaper* via the UAS element, which is expected to impair or ablate IPCs. In the absence of RU486, GeneSwitch-GAL4 is expected to remain inactive (**Figure 1C**).

### PCR genotyping and sanger sequencing

Genomic DNA was isolated from a single female fly per line, using a Quick-DNA™ Miniprep Plus Kit (Zymo Research, D4068S) according to the manufacturer’s instructions. FASTA sequences for the *dilp2* and *reaper* genes were retrieved from FlyBase, and primers were designed using UniPro UGENE (**Supplementary Table S3**). PCR reactions were performed using OneTaq® DNA Polymerase (NEB, M0480) and Deoxynucleotide (dNTP) Solution Mix (NEB, N0447S) according to the manufacturer’s instructions. Amplified products (328 bp for *dilp2*GS-GAL4 and 805 bp for UAS-*rpr)* were resolved on 1.2% agarose gels. A DNA ladder was used as a reference (VWR® DNA Molecular Weight Markers, 100 bp Ladder, 97063-488). Thermal cycling conditions were as follows: initial denaturation at 94□°C for 3 min; 35 cycles of 94□°C for 45 s, 54□°C for 45 s, and 72□°C for 100 s; followed by a final extension at 72□°C for 7 min. The PCR products were confirmed by Sanger sequencing (Eurofins Genomics). PCR and Sanger sequencing were used to confirm the presence and identity of both transgenic elements in the candidate line.

### Adult RU486 induction and experimental diet

Under experimental conditions, 2- to 4-day-old adult flies were maintained on the yeast–sucrose experimental diet containing 9% sucrose to control dietary sugar levels while allowing mating (**Supplementary Table S2 Experimental Diet**). Flies were then separated by sex and transferred to fresh medium supplemented with either RU486 or matched vehicle. RU486 (mifepristone; 98% purity; Thermo Scientific Chemicals, TS45998-0050) was dissolved in 80% ethanol and added to food at a final concentration of 200 µM. Vehicle food received the same volume of 80% ethanol without RU486. Flies were transferred to fresh vials every other day. Unless otherwise specified, treatments were carried out for 8 days.

### Treatment Duration Optimization

To identify an induction duration suitable for downstream metabolic assays, 2-day-old female *dilp2*GS*-rpr* and matched control flies were separately transferred to food supplemented with either RU486 or vehicle and maintained for 8, 10, 12, or 14 days. Flies were subsequently collected for glucose assay as described below. For each line, treatment, and time point, three biological replicates were included, with each replicate comprising five flies.

### Food Intake Assay

Food intake was measured using a blue dye feeding assay based on a previously published protocol (Pereira et al., 2018), with minor modifications. 2- to 4-day old flies were maintained on a standard 9% sucrose experimental diet for 2 days (**Supplementary Table S2 Experimental Diet**). Female and male flies were then separated and transferred to standard food supplemented with either 200 µM RU486 and 2% FD&C Blue Dye #1 (OliveNation) or vehicle with 2% FD&C Blue Dye #1, applied to the food surface. A total of 50 µL of dye mixture was added to each vial, and adult flies were allowed to feed for 3 hours during the same time window for all groups. After feeding, flies were anesthetized and inspected under a stereomicroscope. Flies with visible dye on the external body surface were excluded to avoid surface-contamination artifacts. Non-contaminated flies were collected in groups of four and homogenized in 200 µL 1× PBS. Absorbance was measured at 630 nm, and dye concentration was calculated from a standard curve and normalized to body weight.

### Quantitative RT-PCR

To test whether RU486 induction of the constructed *dilp2*GS*-rpr* donor line reduced IPC-associated *dilp2* expression, we measured *dilp2* mRNA by qRT-PCR in adult female *dilp2*GS*-rpr* flies treated with RU486 or vehicle. 2- to 4-day-old adult female *dilp2*GS*-rpr* flies were transferred to experimental food supplemented with either 200 µM RU486 or matched vehicle, as described above, and maintained for 8 days. Total RNA was extracted from whole-body samples consisting of five female flies per biological replicate, with three biological replicates per treatment group, using a Quick-RNA Miniprep Kit (Zymo Research, R1055), according to the manufacturer’s instructions.

First-strand cDNA synthesis was performed using SuperScript™ II reverse transcriptase (Invitrogen, 18064022). For qPCR, 500 ng of cDNA template was used per 20 µL reaction with igScript™ SYBR Green qPCR 2X Master Mix (Intact Genomics, 3356) following the manufacturers’ instructions. qPCR was performed on a CFX96 Touch Real-Time Detection System (Bio-Rad). Gene expression levels were normalized to the reference gene *Actin5C*. Primers for *Actin5C* and *dilp2* were used as previously described (Ismail et al., 2015). Thermal cycling conditions were: 95□°C for 15 min, followed by 35 cycles of 95□°C for 5 s and 54□°C for 30 s. Relative expression was calculated using the 2^−ΔΔ Ct^ method, with vehicle-treated *dilp2*GS*-rpr* flies used as the reference group.

### Glucose, glycogen, and triglyceride assays

Metabolic assays were performed using a modified version of a previously published protocol (Tennessen et al., 2014). Whole-body homogenates were prepared from pre-weighed groups of five same-sex flies in 100 µL of ice-cold 1× PBST (0.1% Tween-20). For glucose and glycogen measurements, 5 µL of homogenate per sample was analyzed in triplicate. Free glucose and total glucose after amyloglucosidase (Sigma, A1602-25MG) digestion were measured using the Infinity Glucose Hexokinase Reagent (Thermo Scientific, TR15421). Absorbance was measured at 340 nm using an Epoch2 plate reader. Glycogen content was calculated as the difference between total and free glucose. For triglyceride measurements, homogenates were diluted 1:3, and 2 µL of the diluted homogenate was used for analysis with the Infinity™ Triglycerides Reagent (Thermo Scientific, TR22421) according to the manufacturer’s instructions. Absorbance was measured at 540 nm using an Epoch 2 plate reader. Metabolite values were normalized to body weight. Each biological replicate consisted of five flies, and each condition included three biological replicates.

### Statistical analysis

Statistical analyses and graph generation were performed in R. For all analyses, fly line was defined as *dilp2*GS*-rpr* or matched control, and treatment was defined as RU486 or vehicle. Continuous outcomes were analyzed using analysis of variance (ANOVA) or equivalent linear model-based approaches appropriate for experimental design. When applicable, models included genotype, treatment, sex, treatment duration, or experimental group as factors. Pairwise comparisons were performed between RU486-treated flies and the corresponding vehicle-treated controls within each line.

For treatment-duration optimization, glucose levels were analyzed separately at each treatment duration using a linear model including line, treatment, and the line × treatment interaction. Planned contrasts compared RU486-treated and vehicle-treated flies within each line. The line × treatment interaction was used to determine whether the RU486 response differed between the *dilp2*GS-*rpr* recombinant line and the matched background control line.

Food intake data were analyzed separately for female and male flies, with line, treatment, and line-by-treatment interaction included in the model. Planned contrasts compared RU486-treated and vehicle-treated flies within each line.

For metabolic phenotype analyses, only female flies were included. Glucose, glycogen, and triglyceride levels were compared among experimental groups. For each phenotype, data were analyzed using a linear model including line, treatment, and the line × treatment interaction. Planned contrasts compared the induced condition, *dilp2*GS*-rpr* + RU486, with the three control conditions: *dilp2*GS*-rpr* + vehicle, matched control + RU486, and matched control + vehicle. Additional planned contrasts compared RU486 and vehicle within each line.

Data are presented as mean ± SEM. Statistical significance was defined as *P* < 0.05. Significance levels are indicated as follows: \**P* < 0.05, \*\**P* < 0.01, \*\*\**P* < 0.001, and n.s., not significant.

## Results

### Construction and genetic validation of the *dilp2*GS*-rpr* recombinant donor line

PCR genotyping was performed to validate the identity of the candidate recombinant chromosome recovered from female meiosis. Amplification of transgene-specific fragments revealed that the *dilp2*GS-GAL4-specific band (∼328 bp) was detected in the *dilp2*GS-GAL4 parental line (lane 2), while the UAS-*rpr*-specific band (∼805 bp) was detected in the UAS-*rpr* parental line (lane 3) (**Figure 1D**). Notably, both bands were simultaneously present in the candidate *dilp2*GS*-rpr* recombinant fly (lane 4), confirming the presence of both *dilp2*GS-GAL4 and UAS-*rpr* transgenes (**Figure 1D**). Sanger sequencing further verified the identity of the amplified products. Together with the recombination scheme and the stable recovery of the recombinant chromosome over the *CyO* balancer, these results support the establishment of the *dilp2*GS-*rpr* recombinant donor line.

### Selection of an 8-day RU486 induction condition based on glucose response

To determine an appropriate treatment duration for inducing metabolic phenotypes after adult-specific IPC ablation, glucose levels were measured in female flies after 8, 10, 12, or 14 days of RU486 treatment. RU486-treated *dilp2*GS*-rpr* flies showed a significant increase in whole-body glucose levels compared with vehicle-treated transgenic controls as early as day 8 (**Figure 2A–E**). This difference remained evident across the later treatment durations, indicating that 8 days of RU486 exposure was sufficient to produce a detectable glucose phenotype in the *dilp2*GS*-rpr* line. In contrast, RU486 treatment did not produce a comparable increase in glucose levels in the matched control line, supporting that the observed glucose phenotype was primarily associated with RU486-induced activation of the *dilp2*GS-*rpr* recombinant chromosome rather than a nonspecific effect of RU486 exposure. Based on these results, an 8-day treatment period was selected for subsequent metabolic analyses.

**Figure 2.**
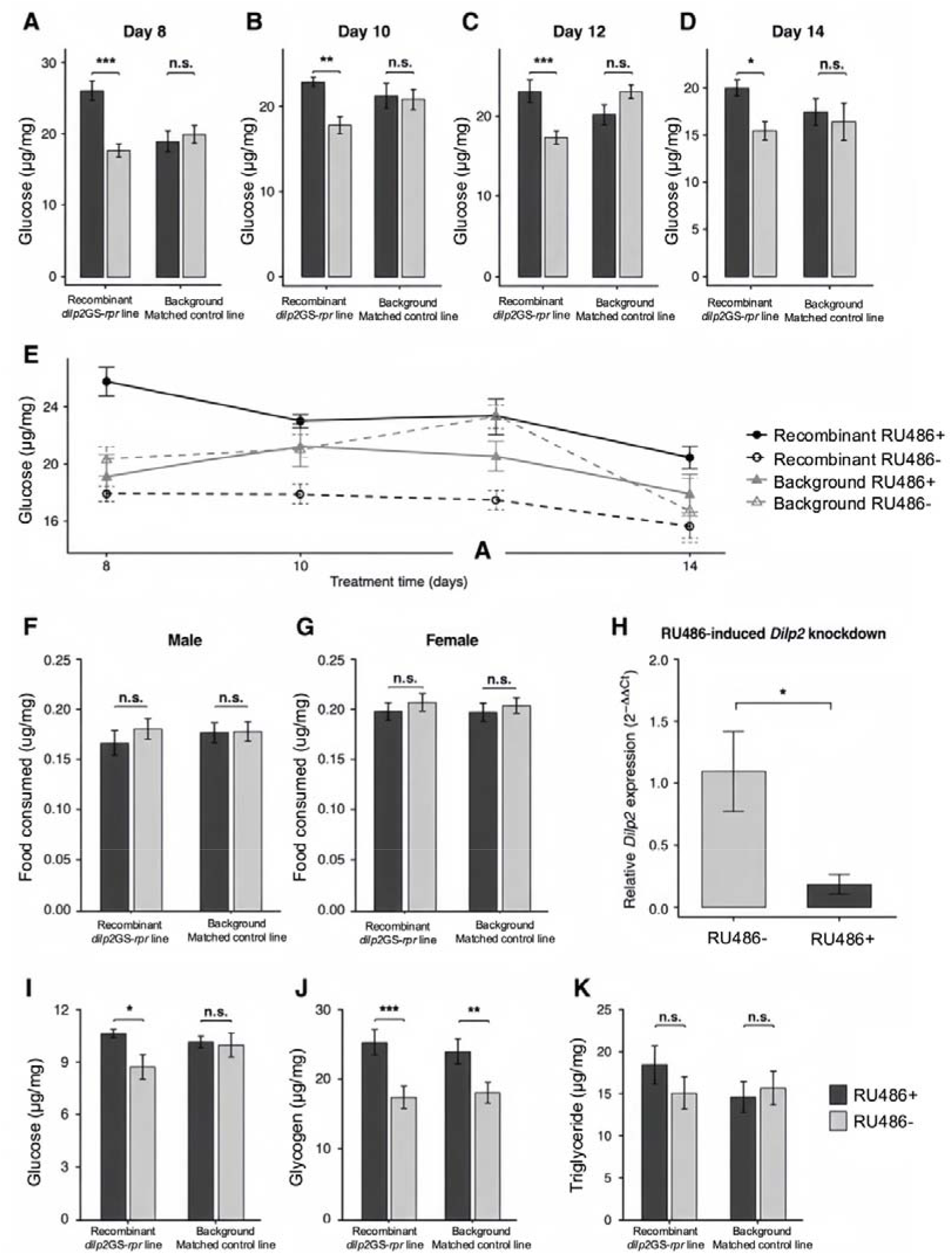
RU486 treatment, food intake, *dilp2* expression, and metabolic measurements in recombinant and background control flies. (A–D) Whole-body glucose levels in female recombinant *dilp2*GS*-rpr* line and the background-matched control line after 8, 10, 12, or 14 days of RU486 or vehicle treatment. (E) Glucose levels across the 8-, 10-, 12-, and 14-day treatment time course in the recombinant *dilp2*GS*-rpr* line and the background-matched control line treated with RU486 or vehicle. (F, G) Short-term dye-based food intake measured by blue dye feeding assay in male (F) and female (G) recombinant *dilp2*GS*-rpr* line and the background-matched control line treated with RU486 or vehicle. (H) Relative *dilp2* mRNA expression measured by qPCR in adult female recombinant *dilp2*GS*-rpr* flies treated with RU486 or vehicle. Expression was normalized to *Actin5C* and calculated using the 2^-ΔΔCt^ method. (I–K) Whole-body glucose (I), glycogen (J), and triglyceride (K) levels in female flies from the recombinant *dilp2*GS*-rpr* line and the background-matched control line after 8 days of RU486 or vehicle treatment. Data are presented as mean ± SEM. RU+, RU486-treated; RU−, vehicle-treated. Statistical comparisons shown above bars compare RU486-treated flies and the corresponding vehicle-treated controls within each line. \**P* < 0.05, \*\**P* < 0.01, \*\*\**P* < 0.001; n.s., not significant.

### RU486 does not detectably alter short-term dye-based food intake

Because altered feeding could confound interpretation of metabolic phenotypes, short-term dye-based food intake was compared between RU486-treated and vehicle-treated flies within each line and sex. No significant difference in food intake was observed between RU486-treated and vehicle-treated groups in either male or female flies for either the *dilp2*GS*-rpr* line or the matched control line (**Figure 2F, G**). These results indicate that RU486 supplementation with did not detectably alter acute dye-based food intake under the experimental conditions used in this study. Therefore, the metabolic changes observed after RU486 treatment are unlikely to be explained by large differences in food intake.

### RU486 treatment reduces IPC-associated *dilp2* expression in the *dilp2*GS*-rpr* line

To test whether RU486 induction of the constructed *dilp2*GS-*rpr* line reduced IPC-associated *dilp2* expression, we quantified *dilp2* mRNA by qPCR after RU486 or vehicle treatment. RU486-treated *dilp2*GS*-rpr* flies showed a significant reduction in *dilp2* mRNA levels compared with vehicle-treated *dilp2*GS*-rpr* controls (**Figure 2H**). Given that *dilp2* is specifically expressed in IPCs, this reduction supports RU486-dependent impairment or partial ablation of IPCs, as indicated by decreased IPC-associated *dilp2* expression. Together with the food intake assay, this result supported the use of the 8-day RU486/vehicle treatment condition for subsequent metabolic analyses.

### Glucose provides the strongest functional metabolic validation of *dilp2*GS*-rpr* induction

To determine whether RU486-induced activation of the recombinant *dilp2*GS*-rpr* line produced metabolic consequences of IPC ablation, we measured whole-body glucose, glycogen, and triglyceride levels after 8 days of RU486 treatment. The main statistical model tested the effects of line, treatment, and line × treatment interaction, allowing us to determine whether the RU486 response differed between the recombinant *dilp2*GS-*rpr* line and the matched background control line (**Supplementary Table S4**).

The clearest line-specific effect was observed for glucose. RU486-treated *dilp2*GS*-rpr* flies showed a significant increase in glucose compared with vehicle-treated *dilp2*GS*-rpr* controls, whereas the background-matched control genotype showed no corresponding glucose change (**Figure 2I, Supplementary Table S5**). This line-specific glucose response indicates that RU486 induction of the *dilp2*GS*-rpr* recombinant chromosome produced a hyperglycemic phenotype, in line with previous studies showing that reduced IPC/DILP function causes fasting hyperglycemia, impaired glucose clearance, or elevated carbohydrate levels (Broughton et al., 2005; Haselton et al., 2010; Rulifson et al., 2002).

We also assessed glycogen and triglyceride levels to determine whether the glucose phenotype was accompanied by broader changes in energy storage. Glycogen increased after RU486 treatment in the *dilp2*GS*-rpr* recombinant line, a direction previously reported after adult IPC ablation (Broughton et al., 2005; Haselton et al., 2010). However, the matched control line also showed a significant RU486-associated glycogen increase (**Figure 2J, Supplementary Table S5**), and the two-factor analysis detected a significant treatment effect for glycogen (**Supplementary Table S4**). This pattern indicates glycogen treatment-sensitive but not specific to the *dilp2*GS*-rpr* line under the current conditions. Thus, although the glycogen increase in RU486-treated *dilp2*GS*-rpr* flies is compatible with reduced IPC/DILP activity (Haselton et al., 2010), the parallel increase in the matched control suggests that RU486 exposure also contributed to glycogen accumulation.

In contrast, whole-body triglyceride levels were not significantly altered by RU486 treatment in either line (**Figure 2K, Supplementary Table S4**). Although prior IPC-ablation studies reported increased stored triglyceride, circulating triglyceride, or lipid storage after reduced IPC/DILP function (Broughton et al., 2005; Haselton et al., 2010), this response was not detected under our 8-day induction condition. Thus, among the metabolic traits tested, whole-body glucose showed the clearest line-specific response to *dilp2*GS*-rpr* induction, whereas glycogen was treatment-sensitive across both lines and triglyceride did not provide a robust validation phenotype under the current conditions.

## Discussion

In this study, we generated and validated the *dilp2*GS*-rpr* recombinant donor line, an adult-inducible IPC-ablation resource carrying *dilp2*GS*-GAL4* and *UAS-rpr* in cis on the same second-chromosome homolog. This design combines IPC-targeted GeneSwitch control with RU486-dependent induction, allowing post-developmental IPC perturbation while keeping the driver and responder linked as one balanced, inheritable donor chromosome. PCR genotyping and sequencing confirmed the presence of both component sequences within the *dilp2*GS-*rpr* recombinant candidate. RU486 treatment reduced *dilp2* mRNA expression, supporting RU486-dependent IPC ablation and providing molecular evidence that the constructed donor line responds to induction as expected.

Metabolic validation further showed that RU486 induction of the constructed *dilp2*GS*-rpr* line produced the expected functional consequences of adult IPC ablation. Whole-body glucose was the strongest functional phenotype: RU486 increased glucose in the *dilp2*GS*-rpr* line but not in the matched control line, consistent with the established role of IPCs and DILP signaling in carbohydrate homeostasis. Previous IPC/DILP perturbation studies have shown elevated carbohydrate levels and impaired glucose clearance (Broughton et al., 2005; Haselton et al., 2010; Rulifson et al., 2002). The matched-control design also clarified how additional metabolic traits should be used in future screens: glycogen appeared sensitive to RU486-associated conditions, whereas triglyceride was not detectably altered under the current induction condition. These results support prioritizing glucose as the primary validated screening phenotype, while treating glycogen and triglyceride as secondary phenotypes that may require further optimization of induction duration, diet composition, or normalization strategy.

A major advantage of the *dilp2*GS-*rpr* recombinant line is its portability: the driver and responder are linked in cis on one balanced donor chromosome, allowing the same inducible IPC-perturbation module to be crossed into diverse recipient backgrounds. This design simplifies modifier-mapping crosses by reducing the need to track independent GAL4 and UAS elements while preserving the perturbation as a fixed donor genotype. The donor chromosome can be introduced into homozygous inbred panels, recombinant inbred panels, or other genetically defined backgrounds, including the DGRP, DSPR (King et al., 2012), Global Diversity Lines (Grenier et al., 2015), African or European population-derived lines (Lack et al., 2015), or laboratory mutant backgrounds to test how recipient genetic variation modifies RU486-induced IPC/DILP phenotypes. Similar donor/tester strategies have been used in *Drosophila* genetic-background screens, including linked GAL4/UAS tester chromosomes and multitransgenic, RNAi-based, or promoter-driven donor stocks. In these studies, standardized perturbation stocks were crossed into stable recipient backgrounds to identify natural modifiers of metabolic, degenerative, and other disease-relevant phenotypes (Chow et al., 2016; He et al., 2014; Palu et al., 2019; Palu et al., 2022; Park et al., 2014; Yang et al., 2023).

This resource provides both a validated adult IPC/DILP perturbation line and a practical donor-line framework for modifier mapping. Future applications can prioritize glucose as the primary validated metabolic readout while testing whether additional conditions reveal robust background-dependent effects on glycogen, triglyceride, fecundity, lifespan, body weight, or transcriptomic responses. Additional validation, such as direct visualization of IPC loss would further strengthen the model before large-scale screening and help distinguish true modifier effects from background-dependent differences in GeneSwitch or GAL4/UAS activity (Yang et al., 2023). More broadly, the construction and validation strategy used here can guide the development of similar linked donor-line resources for other inducible transgenic perturbations, enabling defined genetic perturbations to be tested reproducibly across diverse *Drosophila* backgrounds.

## Acknowledgements

We thank Heinrich Jasper for providing the *dilp2-GeneSwitch-*GAL4*/CyO* driver line, and Martin Kreitman and Michael Ludwig for helpful discussions and insights. We also acknowledge the Bloomington *Drosophila* Stock Center for providing fly stocks used in this study. This work was supported by the National Institute of General Medical Sciences of the National Institutes of Health under award numbers R15GM152956 and R35GM160135 to X.Z. The content is solely the responsibility of the authors and does not necessarily represent the official views of the National Institutes of Health.

## Author contributions

X.Z. conceived the study. Y.C. generated and validated the recombinant *dilp2*GS-*rpr* line, performed the experimental assays, interpreted food intake results, generated figures and tables, and contributed to manuscript writing and revision. Y.B. performed the metabolic and statistical analyses, generated figures and tables, interpreted the metabolic results, and contributed to manuscript writing and revision. X.Z. provided project planning, supervision, funding support, and critical manuscript revision. All authors reviewed and approved the final manuscript.

## Data availability

The authors declare that the main data supporting the findings of this study are available within the article and its Supplementary Information files. Extra data are available from the corresponding author upon request.

## Appendix A Supplementary Data

**Supplementary Table S1**. Genotypes of the parental fly lines and *dilp2*GS-*rpr* donor line.

**Supplementary Table S2**. Experimental Diet and Standard Maintenance Diet.

**Supplementary Table S3**. PCR and q-RT-PCR primer sequences.

**Supplementary Table S4**. Two-way ANOVA of line and RU486 treatment effects on metabolic phenotypes.

**Supplementary Table S5**. Within-line RU486 treatment contrasts for metabolic phenotypes.

